# Drug susceptibility profiling of Australian *Burkholderia* species as models for developing melioidosis therapeutics

**DOI:** 10.1101/2020.01.21.914846

**Authors:** Anna S. Amiss, Jessica R. Webb, Mark Mayo, Bart J. Currie, David J. Craik, Sónia Troeira Henriques, Nicole Lawrence

**Affiliations:** Institute for Molecular Bioscience, The University of Queensland, Brisbane, Queensland, 4072, Australia; Global and Tropical Health Division, Menzies School of Health Research, Darwin, Northern Territory, 0811, Australia; Department of Infectious Diseases and Northern Territory Medical Program, Royal Darwin Hospital, Darwin, Northern Territory, 0811, Australia; School of Biomedical Sciences, Institute of Health & Biomedical Innovation, Queensland University of Technology, Translational Research Institute, Brisbane, Queensland, 4102, Australia

## Abstract

**Background:** Melioidosis is a neglected tropical disease caused by the Gram-negative soil bacterium *Burkholderia pseudomallei.* Current treatment regimens are prolonged and costly, and acquired antimicrobial resistance has been reported for all currently used antibiotics.

**Objectives:** Efforts to develop new treatments for melioidosis are hampered by the risks associated with handling pathogenic *B. pseudomallei*, which restricts research to facilities with Biosafety Level (BSL) 3 containment. Closely related *Burkholderia* species that are less pathogenic can be investigated under less stringent BSL 2 containment. We hypothesized that near-neighbour *Burkholderia* species could be used as model organisms for developing therapies that would also be effective against *B. pseudomallei*.

**Methods:** We used microbroth dilution assays to compare the susceptibility of three Australian *B. pseudomallei* isolates and five near-neighbour *Burkholderia* species – *B. humptydooensis, B. thailandensis, B. oklahomensis, B territorii* and *B. stagnalis –* to antibiotics currently used to treat melioidosis, and general-use antibacterial agents. We also established the susceptibility profiles of *B. humptydooensis* and *B. territorii* to 400 compounds from the Medicines for Malaria Venture Pathogen Box.

**Results:** From these comparisons, we observed a high degree of similarity in the susceptibility profiles of *B. pseudomallei* and near-neighbour species *B. humptydooensis, B. thailandensis, B. oklahomensis* and *B. territorii.*

**Conclusions:** Less pathogenic Australian *Burkholderia* species *B. humptydooensis, B. thailandensis, B. oklahomensis* and *B. territorii* are excellent model organisms for developing potential new therapies for melioidosis.

## Introduction

*Burkholderia pseudomallei* is a Gram-negative bacterium that causes melioidosis,^1, 2^ a neglected tropical disease with an estimated 165,000 cases and 89,000 deaths per year.^1^ Mortality rates for infected individuals vary between 10% in Darwin (Northern Territory, Australia),^3^ where state-of-the-art intensive care facilities are available; and over 40% in endemic regions in southeast Asia, where health resources are more limited.^4^ *B. pseudomallei* is intrinsically resistant to many antibiotics, which limits treatment options; but importantly, environmental isolates and primary *B. pseudomallei* isolates (from melioidosis patients prior to antibiotic exposure) are almost universally susceptible to the first-line drugs used for melioidosis therapy, including ceftazidime, meropenem and cotrimoxazole.^5-7^

In instances where melioidosis is incorrectly diagnosed, initial treatment includes conventional large spectrum antibiotic classes, such as aminoglycosides (e.g. streptomycin, gentamicin and kanamycin), early generation β-lactam antibiotics (e.g. penicillin), fluoroquinolones (e.g. ciprofloxacin) and macrolides (e.g. erythromycin). These generalised treatments have little effect on *B. pseudomallei*, and therefore, result in a low rate of success during periods of misdiagnosis.^6, 8-10^ The limited effectiveness of many therapeutics against *B. pseudomallei* is due to intrinsic and developed resistant to many antibiotics, via a number of different mechanisms including reduced permeation,^11^ drug efflux,^5, 12, 13^ enzymatic drug inactivation,^14, 15^ or mutations.^16-18^

The current therapeutic strategy for treating correctly diagnosed melioidosis patients involves a two-phase schedule, comprising an acute intravenous phase followed by an oral eradication phase.^19^ The standard first-line therapy in Australia for the acute phase is ceftazidime for 10 – 14 days, while meropenem is used for severe infections or where treatment with ceftazidime has failed.^6, 19^ The length of the oral eradication phase, which is most commonly co-trimoxazole (trimethoprim-sulfamethoxazole), is correlated with the success of treatment and reduction in frequency of relapse, often lasting four to six months.^6, 19, 20^ The prolonged nature of the melioidosis treatment schedule can lead to acquired resistance, a significant event that has been linked to treatment failure and mortality in melioidosis patients from the Northern Territory.^5^ Prolonged and costly treatments are especially undesirable in many regions where melioidosis is endemic,^1, 2, 8^ and to overcome both intrinsic and acquired antibiotic resistance, more efficacious therapeutics for the treatment of melioidosis are required.^1, 21^

Efforts to develop new treatments for melioidosis are hampered by the classification of *B. pseudomallei* as a risk group 3 microorganism (i.e. the potential to cause serious human disease) in most countries.^22-25^ This classification restricts its research in laboratories classified as biosafety level 3 (BSL 3 in United States of America^26^ or the equivalent physical containment (PC) 3 in Australia and New Zealand^27^). *B. pseudomallei* is also recognised as a tier 1 biothreat agent on the Centre for Disease Control and Prevention Bioterrorism Agent list,^25^ a classification that further restricts research efforts.^28, 29^ To address this restriction, mutant *B. pseudomallei* strains Bp82 and Bp190, were produced as laboratory models that are avirulent to mice and hamsters.^30^ However, naturally occurring *Burkholderia* species that are not implicated in human disease, have also been described in terms of their close relatedness to *B. pseudomallei*,^31-33^ and these species warrant further investigation as model organisms.

### Objectives

With the aim of overcoming the limitations of containment and handling restrictions of *B. pseudomallei*, we set out to characterise the antibiotic susceptibility profiles of closely related but non-pathogenic *Burkholderia* species, and establish safer model organisms for melioidosis research that can be conducted in BSL 2 facilities. On the basis of genetic relatedness, *B. thailandensis* has previously been used as model for *B. pseudomallei*,^34-36^ but comparison of its antibiotic susceptibility has not been extensively evaluated. Therefore, we included *B. thailandensis* along with representative species – *B. humptydooensis, B. oklahomensis, B territorii* and *B. stagnalis*– in our susceptibility profiling studies.

## Methods

### *Burkholderia* isolates

*B. pseudomallei* and near-neighbour isolates were collected from environmental samples (Menzies School of Health Research) using previously developed methods.^37-39^ *Burkholderia* isolates used in this study include *B. pseudomallei* (MSHR10517, MSHR2154 and MSHR1364), *B. humptydooensis* (MSMB043), *B. oklahomensis* (MSMB0175), *B. stagnalis* (MSMB049), B. *thailandensis* (MSMB0608) and *B. territorii* (MSMB0110).

### Antibiotic panel

Antibiotics were selected to represent the current standard therapeutics for treating melioidosis, ceftazidime, co-trimoxazole and meropenem;^6^ and antibiotics more generally used in a clinical setting for treating bacterial infections, such as doxycycline and amoxicillin. To allow a broader characterisation, additional antibiotics with varying levels of efficacy against *B. pseudomallei* ^9, 40-46^ were also included. A comprehensive overview of the therapeutic target, mode of action, and expected dose required to inhibit 90% or 100% of *B. pseudomallei* growth is shown for each of the antibiotics in Table S1.

### Antibiotic susceptibility profiles

Antimicrobial susceptibility testing was performed using a plate-based broth microdilution method.^47^ Briefly, assays were performed at 30 °C in Mueller Hinton broth (MHB) with bacteria in mid log phase growth that were diluted to ∼ 10^6^ colony forming units/mL (OD_600_ = 0.001). Compounds were prepared in water or dimethyl sulfoxide (DMSO) and two-fold serial dilutions in MHB were added to the bacteria (final bacterial concentration ∼ 5 × 10^5^ CFU/mL, with a maximum of 0.64% (v/v) DMSO). The minimal inhibitory concentration (MIC) was determined to be the lowest concentration of compound that inhibited visible bacterial growth 24 h after treatment. Resazurin (final concentration 0.001% (w/v)) was added to each well for an additional one hour to confirm MIC visualisation. Resazurin (blue) is an oxidation-reduction indicator of aerobic and anaerobic respiration and is converted to resorufin (pink) by viable cells. MIC was determined from the well with the lowest compound concentration that remained blue (no respiration).

### Medicines for Malaria Venture (MMV) Pathogen Box compound susceptibility profiles

A Pathogen Box with 400 drug-like compounds was provided by the MMV.^48^ Compounds were supplied as 10 mM stock solutions in 100% DMSO, and were diluted in MHB according to the suggested procedure in the Pathogen Box supporting documentation. Initial antimicrobial susceptibility testing of the 400 compounds was performed at 20 μM, using the broth microdilution method as described above. Ceftazidime (20 μM) was added as a control to each plate as a positive control for 100% growth inhibition.^6^ Subsequently, compounds with observed activity at 20 μM were serially diluted to determine the MIC. Compound ID, molecular weight, molecular formula and structure of compounds with activity against *B. humptydooensis* and *B. territorii* at 20μM are provided in Figure S1.

## Results and Discussion

### *Burkholderia* near-neighbour isolates

*B. pseudomallei* belongs to the genus *Burkholderia*, which comprises over 70 species with varying virulence and pathogenicity.^49-51^ These species are divided according to their close relationship to either *B. pseudomallei* (the *B. pseudomallei* complex [Bpc]) or *B. cepacia* (the *B. cepacia* complex [Bpc]). In the current study, we have included *B. pseudomallei* and five representative near-neighbour *Burkholderia* species, *B. thailandensis, B. humptydooensis, B. oklahomensis, B. stagnalis* and *B. territorii*. The relationship between the near-neighbour species and pathogenic *Burkholderia* species is represented in Figure 1. *B. thailandensis, B. humptydooensis* and *B. oklahomensis* are most closely related to *B. pseudomallei* and fall within Bpc,^31, 32, 52^ whereas *B. stagnalis* and *B. territorii* are more closely related to *B. cepacia, B. cenocepacia* and *B. multivorans*) and fall within Bcc.^33, 52^ Genetic distance from *B. pseudomallei* is shown in Table S2 for near-neighbour isolates from a previous study.^52^

**Figure 1.**
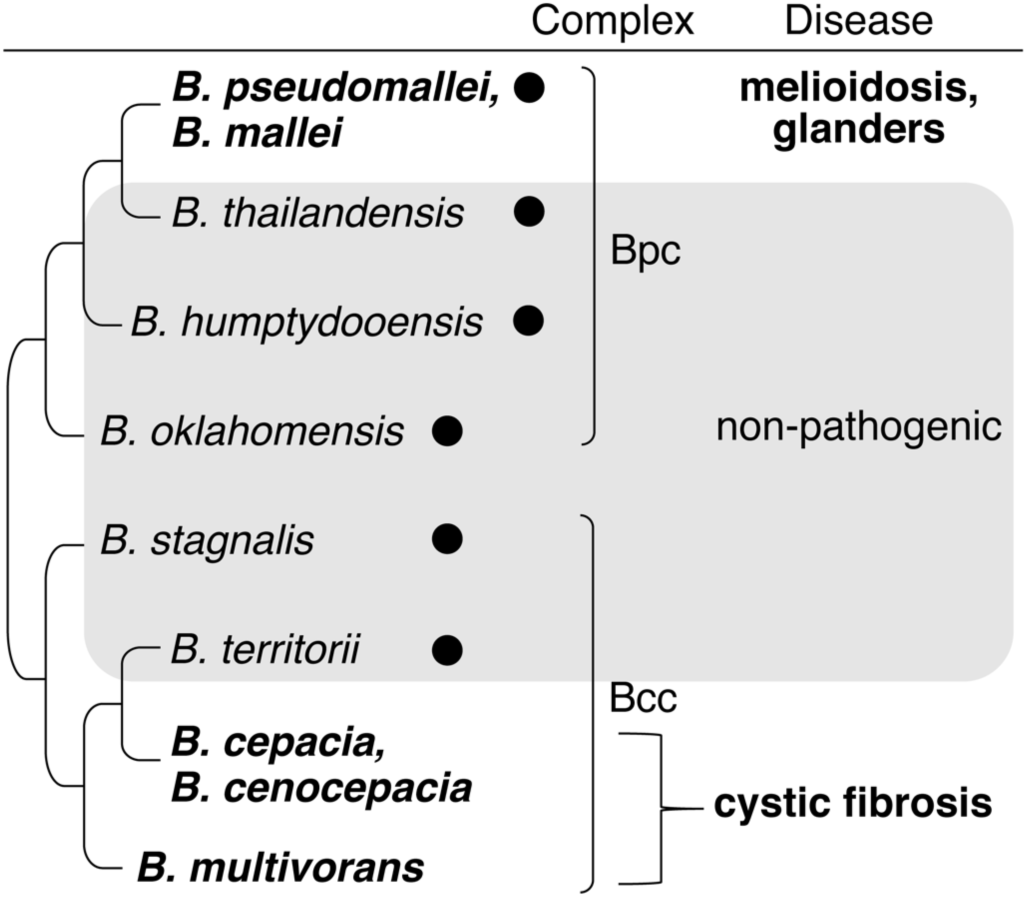
Schematic representation of near-neighbour *Burkholderia* species and their relatedness to *B. pseudomallei* and other major disease-causing species. Relationships are derived from previous phylogenetic trees.^31, 33, 52^ Lines represent relationships between the species, but not genetic distance. Closed circles indicate *Burkholderia* species included in this study. The species highlighted in bold are implicated in human disease.

### Antibiotic susceptibility profiles for *B. pseudomallei* and near-neighbours

The aim of this investigation was to determine whether the susceptibility of *B. pseudomallei* to a range of therapeutics used to treat melioidosis and generalised bacterial infections is recapitulated by near-neighbour species. We hypothesized that species with similar antibiotic susceptibility profiles to *B. pseudomallei* would have utility as less pathogenic models, to facilitate initial screening of new therapeutic molecules without the restrictive physical containment requirements required for working with *B. pseudomallei*.

Current melioidosis treatment involves a regimen of antibiotics, including ceftazidime or meropenem, with or without co-trimoxazole. Therefore, we compared the susceptibility of *B. pseudomallei* near-neighbour species and *B. pseudomallei* isolates to these three key antibiotics. *B. pseudomallei* isolates MSHR10517, MSHR2154 and MSHR1364 were inhibited by 1 – 3 mg/L of ceftazidime, 6 mg/L co-trimoxazole and 1 – 2 mg/L of meropenem (Table 1). These values were consistent with previously reported MICs for other *B. pseudomallei* isolates (see Table S1).^9, 41, 44, 53^ By comparing the susceptibility of *B. pseudomallei* isolates MSHR10517, MSHR2154 and MSHR1364 to the near-neighbour species, we found that the *B. pseudomallei* susceptibility profile was best reflected by *B. humptydooensis*, which was completely inhibited by 1 – 3 mg/L ceftazidime, 6 – 12 mg/L co-trimoxazole and 1 mg/L meropenem. *B. thailandensis* and *B. territorii* also had similar MIC values for ceftazidime and meropenem but were slightly more susceptible to co-trimoxazole (2 – 3 mg/L). In contrast, *B. oklahomensis* was two to four-fold more susceptible to ceftazidime and co-trimoxazole than the *B. pseudomallei* isolates. *B. stagnalis* was two-fold less susceptible to ceftazidime (MIC 5 mg/L) and co-trimoxazole (MIC 6– 12 mg/L), with greater than ten-fold reduced susceptibility to meropenem (MIC of 14 – 38 mg/L). Together, these susceptibility data show that *B. humptydooensis, B. thailandensis* and *B. territorii* best represent the antibiotic susceptibility of *B. pseudomallei* isolates for first- and second-line melioidosis therapies.

**Table 1:** MICs of antibiotics against *Burkholderia* strains ^a^.

|  |  | MIC (mg/L) <sup>b</sup> |  |  |  |  |  |  |  |  |
| --- | --- | --- | --- | --- | --- | --- | --- | --- | --- | --- |
|  | Antibiotic <sup>c</sup> | Mode of action <sup>d</sup> | *B. pseudomallei* |  |  | *B. hu* | *B. ok* | *B. th* | *B. st* | *B. te* |
|  |  |  | MSHR<br>1364 | MSHR<br>2154 | MSHR<br>10517 | MSMB<br>043 | MSMB<br>0175 | MSMB<br>0608 | MSMB<br>049 | MSMB<br>0110 |
| Line 1 | Ceftazidime | CWS | 3 | 1 – 3 | 3 | 1 – 3 | 0.3 | 1 – 3 | 5 | 3 |
|  | Co-trimoxazole | NAS | 6 | 6 | 6 | 6 – 12 | 1 – 6 | 2 – 3 | 6 – 12 | 2 – 3 |
| Line 2 | Meropenem | CWS | 1 | 1 | 2 | 1 | 0.4 – 2 | 2 | 14 – 28 | 2 |
| General-use | Cefsulodin | CWS | - | - | - | >35 | >35 | >35 | >35 | >35 |
|  | Amoxicillin |  | ≥23 | ≥23 | ≥23 | ≥23 | ≥23 | ≥23 | ≥23 | ≥23 |
|  | Ampicillin |  | ≥24 | ≥24 | ≥24 | ≥24 | ≥24 | ≥24 | ≥24 | ≥24 |
|  | Sulfamethoxazole | NAS | - | - | - | 16 – ≥16 | 8 – ≥16 | ≥16 | ≥16 | ≥16 |
|  | Trimethoprim |  | 9 | 5 – 9 | 9 | 5 – 9 | 1 – 9 | 1 | 2 – 5 | 0.6 – 1 |
|  | Rifampicin |  | 13 | 13 | 13 | 13 | 7 | 13 | 26 – 53 | 13 |
|  | Nalidixic Acid | DR | ≥16 | ≥16 | ≥16 | 8 – 16 | >16 | 16 | >16 | 8 |
|  | Doxycycline | PS | 1 | 1 | 1 | ≤ 0.3 – 1 | 0.3 – 1 | 1 | 8 – 16 | 0.01 – 1 |
|  | Tetracycline |  | 2 | 2 | 2 – 4 | 2 – 7 | 1 – 28 | 2 – 7 | 14 – ≥28 | 0.4 – 2 |
|  | Chloramphenicol |  | 21 | 21 | 21 | 3 – 10 | 5 – 10 | 5 – 10 | 10 – 21 | 5 |
|  | Kanamycin |  | ≥37 | ≥37 | ≥37 | 19 – 37 | ≥37 | ≥37 | ≥37 | 37 – ≥37 |
|  | Gentamicin |  | ≥37 | ≥37 | ≥37 | ≥37 | ≥37 | ≥37 | ≥37 | ≥37 |
|  | Puromycin |  | ≥35 | ≥35 | ≥35 | ≥35 | ≥35 | ≥35 | ≥35 | ≥35 |
|  | Spectinomycin |  | ≥32 | ≥32 | ≥32 | ≥32 | ≥32 | ≥32 | ≥32 | ≥32 |
|  | Clarithromycin |  | - | - | - | ≥48 | ≥48 | ≥48 | ≥48 | ≥48 |
|  | Paromomycin |  | - | - | - | ≥40 | ≥40 | ≥40 | ≥40 | ≥40 |
|  | Streptomycin |  | - | - | - | ≥47 | ≥47 | ≥47 | ≥47 | ≥47 |

<sup>a</sup> Near-neighbour species *B. humptydooensis*, *B. hu*; *B. oklahomensis*, *B. ok*; *B. thailandensis*, *B. th*; *B. stagnalis*, *B. st*; *B. territorii*, *B. te*.
<sup>b</sup> MICs were determined using broth microdilution of bacteria in growth phase. Tested antibiotic concentrations in µM were converted to mg/L.
<sup>c</sup> Antibiotics: Line 1 and Line 2, current therapies for treating melioidosis; General-use, used for unknown bacterial infections.
<sup>d</sup> Mode of action as described by DrugBank.<sup>54</sup> Inhibition of: cell wall synthesis (CWS), nucleic acid synthesis (NAS), DNA replication (DR), protein synthesis (PS).

We also compared the activity of antibiotics that are commonly used to treat bacterial infections (see Table 1, general-use). Doxycycline showed potent activity toward *B. pseudomallei* isolates (MIC 1 mg/L) that was consistent with previous reports.^9, 20^ The near-neighbour species *B. humptydooensis, B. thailandensis, B. oklahomensis* and *B. territorii* were even more susceptible to treatment with doxycycline, with MIC values of 0.01 – 1 mg/L. These near-neighbour species were also more susceptible to treatment with trimethoprim (MIC 0.6 – 9 mg/L) and chloramphenicol (MIC 3 – 10 mg/L), compared to *B. pseudomallei* (MIC 5 – 9 mg/L and 21 mg/L respectively).

The activities of rifampicin (MICs 7 – 13 mg/L) and tetracycline (0.4 – 7 mg/L) against *B. humptydooensis, B. thailandensis* and *B. territorii* agreed with their activity against *B. pseudomallei*, with less than two-fold difference in MIC values (see Table 1). Rifampicin had poor activity against *B. pseudomallei* (MIC 8 – 16 mg/L) that was consistent with previous reports,^45^ and in the current study it also showed poor potency against *B. oklahomensis* (MICs 7 mg/L). Tetracycline was less comparable, with MIC against *B. oklahomensis* of up to 28 mg/L. *B. stagnalis* was less susceptible than *B. pseudomallei* to both tetracycline (MICs > seven-fold higher) and rifampicin (MICs > two-fold higher).

For nalidixic acid and kanamycin, complete inhibition of *B. pseudomallei* isolates was not observed following treatment with 16 mg/L or 37 mg/L antibiotic (the highest concentrations tested), but some activity against *B. humptydooensis* and *B. territorii* was observed at these doses. Consistent with previous reports, amoxicillin, ampicillin, clarithromycin gentamicin, puromycin, spectinomycin and streptomycin had no activity against the *B. pseudomallei* isolates,^6, 9, 12, 44, 45^ and these antibiotics also showed no activity against the near-neighbour species at the tested concentrations. We additionally tested antibiotics with limited or no previously reported susceptibilities to *B. pseudomallei* and showed that cefsulodin and paromomycin were not active against any of the *Burkholderia* species at the tested concentrations.

From these comparative antibiotic susceptibility screens, we showed that *B. pseudomallei* near-neighbour species, *B.humptydooensis, B. thailandensis, B. oklahomensis* and *B. territorii*, have similar antibiotic susceptibility profiles to those of *B. pseudomallei* against key melioidosis therapeutics, as well as a number of other antibiotics used for treating bacterial infections. Furthermore, the similarities in antibiotic susceptibility span across multiple modes of action, including inhibition of cell wall synthesis, nucleic acid synthesis, DNA replication, and protein synthesis. Therefore, we propose that these non-pathogenic *Burkholderia* species can be used as model species to screen and identify novel antibiotics, and to predict potency against *B. pseudomallei*.

### Susceptibility of *B. humptydooensis* and *B. territorii* to MMV compounds

To further evaluate the suitability of the near-neighbour *Burkholderia* species as models for predicting the drug susceptibility of *B. pseudomallei*, we examined the susceptibility of *B. humptydooensis* and *B. territorii* to 400 diverse, drug-like molecules from the MMV Pathogen Box.^48^ This box includes compounds with activity against infectious diseases (including tuberculosis, malaria, and African sleeping sickness), that have recently been examined for activity against five *B. pseudomallei* isolates.^55^

We initially screened the MMV compounds for activity against *B. humptydooensis* and *B. territorii* at 20 μM (∼8 – 16 mg/L), and identified four active compounds in agreement with previously reported activity toward *B. pseudomallei* isolates;^55^ doxycycline, levofloxacin, rifampicin, MMV675968, and MMV688271 (see Figure S1 for characteristics of the compounds). An additional compound, MMV67968, showed novel activity toward the near-neighbour species.

Next, we determined the MICs toward *B. humptydooensis* and *B. territorii* for the compounds identified from the initial screen, and for three additional compounds from a previous *B. pseudomallei* susceptibility screen;^55^ auranofin, miltefosine and MMV688179 (see Table 2 and Table S1). The activities of doxycycline (MIC 0.5 – 1 mg/L), levofloxacin (MIC 1 – 6 mg/L), MMV688271 (MIC 4 – 8 mg/L) and ceftazidime (MIC 2 – 4mg/L) against the near-neighbour species were within two-fold of their reported MICs against *B. pseudomallei* (1 – 3 mg/L, 4 – 10 mg/L and 8 – 12 mg/L respectively).^55^ Notably, the MICs for rifampicin, doxycycline and ceftazidime determined from the MMV compound screen (Table 2) were in close agreement with MICs from the antibiotic susceptibility screen (Table 1). The MIC values for auranofin, miltefosine and MMV688179 were at or above the highest concentration tested. These high MICs are consistent with previous reports in *B. pseudomallei*^55^ (see Table 2).

**Table 2:** MICs of MMV compounds against *B. humptydooensis* and *B. territorii*.

| Compound | Mode of Action <sup>c</sup> | MIC (mg/L) <sup>a</sup> |  | MIC (mg/L) <sup>b</sup> |
| --- | --- | --- | --- | --- |
|  |  | *B. humptydooensis*<br>MSMB 43 WGS | *B. territorii*<br>MSMB 110 WGS | *B.pseudomallei* |
| Ceftazidime <sup>d</sup> | CWS | 2 – 4 | 1 – 4 | 3 – 4 |
| Doxycycline | PS | 0.5 | 0.5 – 1 | 1 – 3 |
| Levofloxacin | DR | 1 – 3 | 1 – 6 | 4 – 10 |
| Rifampicin | NAS | 13 | 13 – 26 | 18 – 45 |
| MMV688271 | unknown | 4 – 8 | 4 – 8 | 6 – 12 |
| MMV675968 | unknown | 6 – 12 | 3 – 12 | n.d. <sup>e</sup> |
| Auranofin | unknown | >22 | >22 | 150 |
| Miltefosine | ET | >13 | 3 - >13 | >1600 |
| MMV688179 | unknown | 15 | 15 | 12.5 - >100 |

<sup>a</sup> MIC values for *B. humptydooensis* and *B. territorii* calculated from serial dilutions of the MMV Pathogen Box compounds, starting at 20 µM. Values were converted to mg/L. <sup>b</sup> MIC values determined by Ross et al, 2018 using *B. pseudomallei* isolates K96243 576, NCTC13178, NCTC13179 and MX2013.<sup>55</sup> <sup>c</sup> Mode of action as described by DrugBank<sup>54</sup> – inhibition of: cell wall synthesis (CWS), nucleic acid synthesis (NAS), DNA replication (DR), electron transport (ET), protein synthesis (PS). <sup>d</sup> Ceftazidime was not part of the MMV panel but was included as a positive control with activity toward *Burkholderia* species. <sup>e</sup> MIC toward *B. pseudomallei* was not determined. Inhibitory activity was not detected at 2 µM (0.72 mg/L).<sup>55</sup>

MMV675968 was active against *B. humptydooensis* and *B. territorii*, with MIC between 3 – 12 mg/L, an activity range that is within two-fold of the ‘gold-standard’ melioidosis therapy ceftazidime. Therefore, this newly identified molecule is worthy of further investigation for activity against *B. pseudomallei.*

Overall, comparison of the activity of the 400 tested MMV compounds against *B. humptydooensis* and *B. territorii* provided independent correlation for four of the seven compounds with previously identified activity against *B. pseudomallei*.^55^ Although almost all strains of *B. pseudomallei* tested have intrinsic resistance to gentamicin and streptomycin, there have been rare reports of susceptibility to these antibiotics in isolates from Thailand and Malaysia.^42, 56^ These examples might suggest differences in susceptibility profiles of *B. pseudomallei* isolates originating from different geographic regions; a question we have not directly answered in this study. However, we show that these near-neighbour isolates provide a strong prediction for susceptibility of Australian *B. pseudomallei* isolates, and can also predict the susceptibility of clinical isolates of Mexican, Thai and Australian origin to 400 compounds.^55^

## Conclusions

In this study, we demonstrate similar susceptibility of non-pathogenic *Burkholderia* species *B. humptydooensis, B. oklahomensis, B. thailandensis, B. territorii* and pathogenic *B. pseudomallei* for an extensive panel of antibiotics and drug-like compounds. In particular, the newly characterised species *B. humptydooensis and B. territorii*, and previously described *B. thailandensis*, provided good correlation with *B. pseudomallei* susceptibility. Thus, these near-neighbour species have potential for use in initial investigations and high throughput screening of molecules for melioidosis therapeutic development.

The lower risk-group classification of the near-neighbour species allows expansion of melioidosis research into a wider landscape, where more laboratories have adequate facilities to perform the initial compound discovery. We are hopeful that inclusion of well characterised and non-pathogenic model organisms in melioidosis research will accelerate the development of new treatment options for this neglected tropical disease.

## Acknowledgments

We would like to thank Dr Alysha Elliot at the Institute for Molecular Bioscience, University of Queensland for her critical review of this work, Medicines for Malaria Venture for supplying the Pathogen Box compounds, and Vanessa Rigas for her laboratory support at Menzies School of Health Research.

## Transparency declaration

None to declare.

## Author contributions

A.S.A., B.J.C., S.T.H., and N.L. designed the study;

A.S.A. and J.R.W. performed experiments with support from M.M., S.T.H., and N.L.;

B.J.C., D.J.C., and S.T.H. provided resources and acquired funding;

A.S.A., J.R.W. S.T.H., and N.L. prepared the original draft, which was reviewed and edited by all authors.

## SUPPLEMENTARY DATA

**Table S1.** Antibiotic characteristics and activity toward *Burkholderia pseudomallei*.

|  | Antibiotic | Drug class | Target for inhibition | Mode of action <sup>a</sup> | MIC 90 (mg/L) | MIC 100 (mg/L) |
| --- | --- | --- | --- | --- | --- | --- |
| Line 1 | Ceftazidime | Cephalosporin | Cell wall synthesis | Inhibits penicillin-binding proteins (PBPs) responsible for cell wall synthesis | 2 – 4<sup>1,2</sup> | 1 – 4<sup>3,4</sup> |
|  | Co-trimoxazole | Mixed | Nucleic acid synthesis | Trimoxazole and sulfamethoxazole mode of action |  | 0.125 – 4<sup>4</sup> |
| Line 2 | Meropenem | Carbapenem | Cell wall synthesis | Binds PBP and inhibits bacterial cell wall synthesis | 1 – 4<sup>3</sup> | 0.5 – 4<sup>3,4</sup> |
| General-use | Cefsulodin | Cephalosporin | Cell wall synthesis | Binds PBPs and inhibits peptidoglycan cross linking | >128<sup>2</sup> |  |
|  | Amoxicillin | Penicillin | Cell wall synthesis | Binds PBPs and inhibits peptidoglycan polymer chain cross linkage | >128<sup>2,5</sup> |  |
|  | Ampicillin | Penicillin | Cell wall synthesis | Binds PBP and inhibits bacterial cell wall synthesis |  | 12.5 – 25<sup>6</sup> |
|  | Sulfamethoxazole | Sulfonamide | Nucleic acid synthesis | Interferes with folic acid synthesis | 320<sup>5</sup> |  |
|  | Trimethoprim | DHFR inhibitor | Nucleic acid synthesis | Inhibits dihydrofolate reductase | 64<sup>5</sup> |  |
|  | Nalidixic Acid | Fluoroquinolone | DNA replication | Inhibits the A subunit of bacterial DNA gyrase | 32<sup>2</sup> | >50<sup>6</sup> |
|  | Rifampicin | Rifampicin | DNA-dependent RNA synthesis | Inhibits bacterial DNA-dependent RNA polymerase | 16 – 32<sup>2,5</sup> | 8 – 16<sup>7</sup> |
|  | Doxycycline | Tetracycline | Protein synthesis | Binds to the bacterial 30S ribosomal subunit, blocking aminoacyl tRNA from binding | 1 – 4<sup>2,5</sup> | 0.25 – 3<sup>4,7</sup> |
|  | Tetracycline | Tetracycline | Protein synthesis | Binds to the bacterial 30S ribosomal subunit, blocking aminoacyl tRNA from binding | 0.5–8<sup>1,5</sup> | 1.6 – 3.1<sup>6</sup> |
|  | Gentamicin | Aminoglycoside | Protein synthesis | Binds to 16S rRNA and protein S12 and interferes with mRNA reading | 64–128<sup>2,5</sup> | > 50<sup>6</sup> |
|  | Kanamycin | Aminoglycoside | Protein synthesis | Binds to 16S rRNA and protein S12 and interferes with mRNA reading | 64<sup>5</sup> |  |
|  | Paromomycin | Aminoglycoside | Protein synthesis | Binds to 16S ribosomal RNA causing defective polypeptide chain production |  | >50<sup>6</sup> |
|  | Spectinomycin | Aminoglycoside | Protein synthesis | Binds to the 30S subunit of the bacterial ribosome. Interferes with initiation of protein synthesis and elongation |  | >64<sup>8</sup> |
|  | Streptomycin | Aminoglycoside | Protein synthesis | Binds to 16S rRNA and protein S12 and interferes with the assembly of initiation complex between mRNA and the bacterial ribosome |  | >50<sup>6</sup> |
|  | Clarithromycin | Macrolide | Protein synthesis | Binds to the 50S ribosomal subunit and inhibits RNA-dependent protein synthesis |  | 32 – 64<sup>7</sup> |
|  | Chloramphenicol | Amphenicol | Protein synthesis | Inhibits peptidyl transferase activity of the bacterial ribosome | 16–32<sup>2,5</sup> | ~6 – 20<sup>6</sup> |

<sup>a</sup> Mode of action as described by DrugBank. <sup>9</sup>

**Table S2.** Comparison of genetic distance between *Burkholderia pseudomallei* and near-neighbour species.

| ***Burkholderia***<br>**near-neighbours** | Changes/site between<br>near-neighbour and<br>*B. pseudomallei* <sup>a</sup> |
| --- | --- |
| *B. thailandensis* | ~ 0.032 |
| *B. humptydooensis* | ~ 0.041 |
| *B. oklahomensis* | ~ 0.054 |
| *B. stagnalis* | ~ 0.094 |
| *B. territorii* | ~ 0.115 |

<sup>a</sup> branch distances (in cm) were determined from a maximum-likelihood phylogeny (Figure 1 from Sahl et al 2016 <sup>10</sup>) and converted to changes/site using the scale bar: 1cm of branch length = 0.01 changes/site.

**Figure S1.**
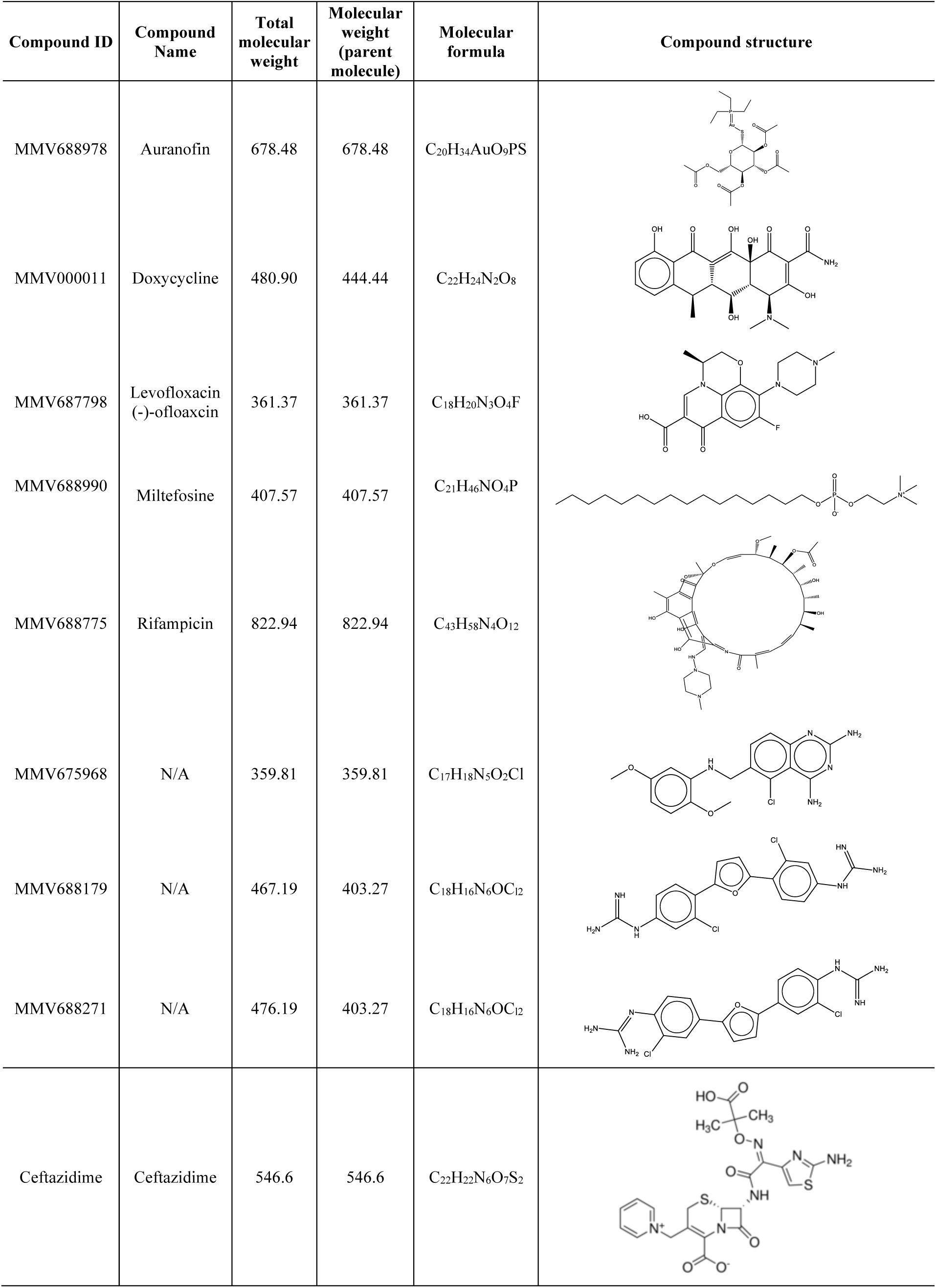
Compounds from the Medicines for Malaria Venture Pathogen Box^11^ with activity against *B. humptydooensis* and *B. territorii* at 20 μM (7.2 – 17.6 mg/L)

